# Androgen Receptor and ACSL4 Interplay as Therapeutic Targets in Breast Cancer

**DOI:** 10.64898/2026.11.21.766272

**Authors:** Melina A. Dattilo, Paula F. Lopez, Yanina Benzo, Hernán Suarez Ceva, Adrián P. Mansini, Cristina Paz, Luke H. Hoeppner, Ernesto J. Podestá, Paula M. Maloberti

## Abstract

High Acyl-CoA synthetase 4 (ACSL4) expression is implicated in the development and progression of aggressive tumors, poor prognosis, and unfavorable clinical outcomes, as observed in triple-negative breast cancer (TNBC). The androgen receptor (AR) has emerged as a therapeutic target in AR-positive TNBCs, while treatment options are even more limited for quadruple-negative breast cancer (QNBC), the AR-negative TNBC subtype. This study investigates the reciprocal regulation between AR and ACSL4 and evaluates their potential as therapeutic targets in breast cancer. Analysis of patient datasets and expression profiling in human breast cancer cells revealed an inverse correlation between ACSL4 and AR expression. We show that ectopic AR expression decreases ACSL4 promoter activity, mRNA, and protein levels, whereas AR inhibition increases ACSL4 expression in breast cancer cells. We demonstrate that AR acts as a repressor by binding to an androgen response element on the ACSL4 promoter. Moreover, ectopic ACSL4 expression decreased AR levels, while pharmacological ACSL4 inhibition or silencing increased AR expression, supporting a negative regulatory loop. To investigate whether ACSL4 inhibition sensitizes QNBC cells to antiandrogen treatment, PRGL493 and bicalutamide (ACSL4 and AR inhibitors, respectively) were combined, producing synergistic inhibitory effects on cell proliferation, migration, and invasion. This combination also showed strong suppressive effects in cancer cell extravasation, a key step in the process of cancer metastasis, using a zebrafish xenotransplantation model. These findings show that ACSL4 inhibition sensitizes QNBC cells to antiandrogen therapy due to reciprocal negative regulation between ACSL4 and AR, providing a novel promising strategy for aggressive breast cancer subtypes.

## Introduction

Triple-negative breast cancer (TNBC) represents around 10-20% of all breast cancers and is characterized by poor prognosis and proneness to metastasis and recurrence as compared to other breast cancer subtypes. The lack of estrogen receptor (ER) and progesterone receptor (PR), and the absence of human epidermal growth factor receptor 2 (HER2) amplification severely restricts therapeutic options, as both endocrine therapies and HER2-directed treatments are ineffective. Consequently, identifying and developing novel therapeutic strategies for TNBC remains a critical unmet need (1). A subset of TNBC has been found to express the androgen receptor (AR) (2), which positions AR signaling as a significant therapeutic target in breast cancer pathogenesis (3). Currently, several clinical trials evaluating AR antagonists commonly used in prostate cancer, such as bicalutamide (BICA) and enzalutamide, have demonstrated clinical benefits and good tolerance in patients with AR-positive TNBC, which paves the way for AR inhibitors as potential treatment options in patients with AR-positive TNBC (4–11). Around 70% of TNBCs, however, lack AR expression and constitute highly aggressive tumors with poor survival and frequent chemoresistance. AR-negative TNBC is known as quadruple negative breast cancer (QNBC) and has been recognized as a highly aggressive breast cancer subtype with limited therapeutic strategies available (12,13).

Proteins such as SKP2, EGFR, immune checkpoint molecules, microRNA signatures, engrailed 1 or Acyl-CoA synthetase 4 (ACSL4) are preferentially expressed in QNBC and could therefore be useful in the development of new therapeutic strategies (13–15). ACSL4, an enzyme involved in arachidonic acid metabolism (16), has been shown to play a pivotal role in promoting tumor aggressiveness in hepatocellular, colorectal, prostate, and breast cancer, among others (17–22). ACSL4 overexpression increases cell proliferation, migration, invasion, tumor growth and resistance to chemotherapeutic drugs or hormone treatments in breast and prostate cancer (20–24). An inverse correlation between ER expression and ACSL4 levels has been observed in TNBC(21,23,25,26). It was previously reported that ER is an indirect negative regulator of ACSL4, as ER overexpression results in a decrease in enzyme expression and transcriptional activity in MDA-MB-231 cells (27). This regulation is reciprocal, as ACSL4 overexpression in ER-positive MCF-7 breast cancer cells is associated with a significant reduction in ER mRNA and protein expression (21,25), which unveils the involvement of ACSL4 overexpression in tamoxifen resistance in breast cancer (23). Moreover, the inhibition of ACSL4 in breast cancer cells has proven to reverse the loss of ER expression (21,25), which sensitizes breast cancer cells to tamoxifen(23,24). Increased ACSL4 levels together with decreased AR expression and androgen insensitivity have been reported in breast cancer (2,13,25), and AR has been postulated as a repressor of ACSL4 in prostate cancer models (28). However, the androgenic regulation of ACSL4 in breast cancer cells has not been described. In this context, the present study aimed to study the implications of ACSL4 in resistance to antiandrogen treatment, analyzing the reciprocal regulation between ACSL4 and AR and its possible therapeutic application in breast cancer.

## Materials and methods

### Reagents

Penicillin-streptomycin, trypsin solution, ethylene diamine tetra-acetic acid (EDTA), and Dulbecco’s modified Eagle’s medium (DMEM) were provided by Gibco-Life Technologies Inc. (Gaithersburg, MD, USA). Fetal bovine serum (FBS) was from PAA laboratories GmbH (Pasching, Austria). BICA, tricaine, phenylthiourea (PTU), dihydrotestosterone (DHT), 3-[4,5-dimethylthiazol-2-yl]-2,5-diphenyltetrazolium bromide (MTT), bovine serum albumin (BSA), molecular weight standards for electrophoresis, hydroxymethyl aminomethane (Tris), sodium dodecyl sulfate (SDS), glycine, tetramethylethylenediamine (TEMED), glutamine, and poly-l-lysine were purchased from Sigma Chemical Co. (St. Louis, MO, US). Opti-MEM, Lipofectamine 2000, and PCR primers were purchased from ThermoFisher Scientific (Waltham, MA, USA), Macrogen Inc. (Seoul, South Korea), and Integrated DNA Technologies (Coralville, IA, US). PVDF membranes, reagents (acrylamide, bisacrylamide, Bradford, and others) and electrophoresis equipment (Mini-Protean II and transfer kits) were provided by BioRad Laboratories (Hercules, CA. US). Enhanced chemiluminescence was from GE Healthcare (Buckinghamshire, UK). Sterile and plastic material for tissue culture was from Orange Scientific (Brainel’Alleud, Belgium). All reagents were of the highest grade available.

### Cell lines and culture conditions

Human breast cancer cell lines MDA-MB-231 and MCF-7 were obtained from the Lombardi Comprehensive Cancer Center (Georgetown University Medical Center, Washington, DC, US). BT-20 and MDA-MB-453 cell lines were obtained from the CIQUIBIC Cell Bank (Universidad Nacional de Córdoba, Argentina). All human cell lines were authenticated by ATCC and used within 10 passages. The absence of mycoplasma was tested monthly by polymerase chain reaction (PCR). As previously described (20,21,23,29), we used modified MCF-7 cells overexpressing ACSL4 (Tet-Off system) and MDA-MB-231 cells with ACSL4 knockdown (shRNA) (29), alongside their respective empty-vector controls. BT-20 and MDA-MB-453 cells were cultured in complete DMEM (10% FBS, 1X glutamine, 100 U/mL penicillin, and 10 μg/mL streptomycin), while MDA-MB-231 and MCF-7 cells required high-glucose DMEM. All cells were maintained at 37 °C in a 5% CO_2_ atmosphere. Experiments involving DHT were performed under steroid-free conditions using phenol red-free medium supplemented with 10% charcoal-stripped FBS, as indicated in the literature (30).

### Plasmids, site-directed mutagenesis, and transfections

For AR overexpression experiments, we used the pSG5-ARh vector (31) or the empty pSG5 vector (Mock) as control. “ACSL4 Promoter” is a reporter plasmid that contains a 1.8 kb fragment of the ACSL4 promoter region (–1681 to +125 bp relative to the transcription start site), cloned between the XhoI and HindIII restriction sites within the multiple cloning region of the pNL1.1 empty vector (Promega Corp., Madison, WI, US), upstream of the NanoLuc® reporter gene (27). ACSL4 Promoter was used as the template for site-directed mutagenesis employing the Quik Change Site-Directed Mutagenesis Kit (Agilent Technologies, Santa Clara, CA, US) following the manufactureŕs instructions. For this purpose, we used primers carrying a point mutation, which allowed the introduction of a CxT substitution at position -912 of the ACSL4 human promoter (Fw: 5′-CAGGAATAGGAAGCTAGTT-3′. Rv: 5′-AACTAGCTTCCTATTCCTG-3′), which in turn eliminates an AR element (ARE) present in ACSL4 Promoter reporter plasmid. The construct obtained was labeled “Mut ARE” and sequenced by Macrogen Inc., confirming the corresponding mutated variant. The effect of Mut ARE was assayed against ACSL4 Promoter as control. Transfections were carried out with Opti-MEM and Lipofectamine® 2000 as previously described (27).

### NanoLuc luciferase reporter assays

The transcriptional activity of reporter plasmids was assessed using the Nano-Glo® Luciferase Assay System (Promega Corp.) as previously described (27). Plasmid pRc/CMVi (Life Technologies, Carlsbad, CA, US) expressing the EGFP gene was used for normalization (32). Luminescence and fluorescence counts were measured in an automated plate reader (Synergy HT, BioTek, Winooski, VT, US). NanoLuc luciferase activity was normalized to EGFP fluorescence counts measured in the same equipment and expressed as arbitrary units (33) **Western blots:** Western blots were performed as previously described (20,21,23,24,27,29). Cells were lysed in RIPA buffer. After protein separation via SDS-PAGE and protein transfer, PVDF membranes were incubated with primary antibodies against ACSL4 (RRID: AB_10843105) or GAPDH (RRID: AB_627678) from Santa Cruz Biotechnology (Dallas, TX, US) or against AR (RRID: AB_10691711) from Cell Signaling Technology (Boston, MA, US) used at 1:1000. Secondary goat anti-mouse (RRID: AB_11125547) and anti-rabbit (RRID: AB_11125142) IgG antibodies conjugated to horseradish peroxidase were purchased from Bio-Rad Laboratories and used at 1:5000. Band intensities were quantified using Gel-Pro Analyzer software, and values were normalized to GAPDH expression.

### RNA extraction and qRT-PCR

RNA was extracted and mRNA levels determined as previously described (20,21,23,24,27,29). AR (Fw: 5′-GGCATGGTGAGCAGAGTG-3′, Rv: 5′-GGGTGGAAAGTAATAGTCAATGG-3′) and ACSL4 (Fw: 5′-CCCGCTATCTCCTCAGACAC-3′, Rv: 5′-CAATTTCATGGCATTTGCAG-3′) mRNA levels were assessed by qPCR using the SYBR Green Master Mix reagent kit (Applied Biosystems, Carlsbad, CA, US) and read in a Mic PCR instrument (Molecular Biosystems, San Diego, CA, USA). Gene expression levels were normalized to human cyclophilin (Fw: 5′-TGGCAAGTCCATCTATGGGGA-3′, Rv: 5′-ACTTATTCGAGTTGTCCAACAGTCAGCA-3′). The reaction conditions for AR were: one 2-min cycle at 95 °C, followed by 40 cycles at 95 °C for 15 s, 55 °C for 30 s, and 72 °C for 30 s. The reaction conditions for cyclophilin or ACSL4 were: one 5-min cycle at 95 °C, followed by 40 cycles at 95 °C for 15 s, 60 °C for 30 s, and 72 °C for 30 s. qPCR data were analyzed by calculating the 2^−ΔΔCt^ value (comparative Ct method) for each experimental sample (relative gene expression).

### Immunofluorescence analysis

Protein expression was analyzed by immunocytochemistry as previously described (29). ACSL4 or AR protein expression was detected using anti-ACSL4 (1:100) or anti-AR (1:100), respectively, followed by a cy3 (red)-conjugated anti-mouse secondary antibody (1:400). Controls omitting the primary antibody were included. The nucleus of each cell was visualized through DAPI counterstain. Images were acquired on a Nikon E200 epifluorescence microscope at 40X objective. Images were obtained at equal gain and exposure time, and the integrated density was measured using FIJI software.

### Cell proliferation assays

Cell proliferation was assessed using MTT as previously described (20). Cells were seeded at a density of 4,000 cells/well in 96-well plates with complete medium and incubated overnight in a humidified atmosphere with 5% CO₂. Cells were then serum-starved overnight before treatment with the agents indicated or vehicle in complete medium for the times specified in the figure legends. MTT (2 mg/ml) was subsequently added and, after 2:30 h, absorbance of the formazan crystals dissolved in DMSO was measured at 570 nm using a Synergy HT microplate reader (Biotek).

### Cell migration assays

The Incucyte® Scratch Wound Assay was performed following supplier’s instructions (Sartorius, Bohemia, NY, US). Briefly, 40,000 MDA-MB-231 cells/well were seeded in 96-well plates and incubated at 37 °C in a humidified atmosphere with 5% CO_2_ until 100% confluence. Cells were then incubated in serum-free medium overnight. Wounds were made using the Woundmaker tool provided by the supplier. After two washes with complete medium, treatments or vehicle were applied for the times indicated in figure legends. Photos were taken using the Incucyte® Live-Cell Analysis System (Sartorius) with a 10X magnification. Cell migration was quantified as a percentage of wound closure compared to the cells at 0 h in each group.

### 3D tumor single spheroid cell invasion into surrounding extracellular matrix

The IncuCyte® 3D Spheroid Invasion Assay was performed following supplier’s instructions (Sartorius). Briefly, 5,000 cells/100 µl per well were seeded in a 96-well round-bottom ultra-low attachment plate and centrifuged at 125 g for 10 min at room temperature. Cells were then incubated at 37 °C in a humidified atmosphere with 5% CO_2_ for 48 h. Once spheroids had reached 500 µm in diameter, 10 µl of complete medium containing the corresponding treatment at 20X was added to each well. Subsequently, 90 µl of 25% Matrigel® (Sigma-Aldrich) was added carefully on ice. The plate was incubated at 37 °C for 30 min to allow Matrigel® polymerization. Post-polymerization, 50 μL/well of complete medium containing treatment or vehicle was added at the final concentration as described in the figure legends. The plate was placed in an incubator at 37 °C in a humidified atmosphere with 5% CO_2_ for 5 days. Images were acquired daily using the Incucyte® Live-Cell Analysis System (Sartorius).

### Zebrafish husbandry

Zebrafish (*Danio rerio*) were handled according to IACUC-approved conditions. Fish were maintained at 28 °C on a 14-h light:10-h darkness cycle. Healthy adult *Tg(fli-GFP)* zebrafish were used for egg production. The developing embryos were incubated at 28 °C and maintained in PTU 0.003% w/v to prevent pigmentation.

### Zebrafish model for cancer cell extravasation

A critical step in the process of cancer metastasis is the extravasation of circulating tumor cells through the endothelium of blood vessels to colonize distant organ sites. To evaluate the impact of drug treatments on cancer cell extravasation, we used a well-established transgenic *Tg(fli-GFP)* zebrafish xenograft model of human cancer cell extravasation as previously described (34–38). This model is ideal for tracking tumor cell movement across blood vessels, as these fish express EGFP under the promoter of *fli1*, an endothelial marker. Treated tumor cells were trypsinized, counted, and labeled with Cell Tracker Orange CMTMR (Invitrogen, Waltham, MA, US) according to the manufacturer’s instructions. Cells were resuspended in PBS containing DNase I and heparin, and 200 cells/6 nl were intracardially microinjected into each anesthetized *Tg(fli-GFP)* zebrafish embryo 3 days post fertilization (dpf). Embryos were maintained in embryo water containing PTU 0.003%. Tricaine was used for anesthesia at a concentration of 0.02% and administered by immersion. Embryos showing cancer cells in the bloodstream after injection were maintained at 34 °C for 24 h and then imaged on a Zeiss LSM 900 confocal microscope using 10X or 20X objective to acquire z-stacked images with ZEISS ZEN 3.11 software. All zebrafish studies were conducted following institutionally approved IACUC protocols.

### Statistical Analysis

Results are expressed as mean ± standard deviation of the mean (SD), unless otherwise stated. Statistical significance was assessed using parametric or non-parametric tests, as appropriate. Comparisons between two independent groups were performed using Student’s *t*-test. For multiple comparisons, one-way or two-way analysis of variance (ANOVA) was performed, as appropriate, and Newman-Keuls was used as a post-test. A *p* value<0.05 was considered statistically significant. All analyses were performed using GraphPad Prism software (version 8.4.2; GraphPad Software, San Diego, CA, US). When appropriate, treatment synergy was assessed using the Bliss independence model(39). This model relies on probability theory to describe the behavior of independent events. In this framework, the expected effect of a drug combination is estimated from the individual effects of each compound, assuming the absence of pharmacological interaction between them. The combined response predicted is calculated according to the equation: E(a + b) = E(a) + E(b) − E(a) × E(b), with effect (E) values expressed as percentages. The magnitude of the interaction at each dose combination is further quantified using the excess over Bliss (EOB) score, which is defined as the difference between the inhibitory effect observed and the effect expected. Drug combinations are classified as synergistic when the effect observed experimentally exceeds the predicted value, antagonistic when it is lower, and additive when the effects observed and expected coincide.

## Results

### ACSL4 and AR expression are inversely correlated in breast cancer

We performed bioinformatic analysis using public databases to explore the relationship between ACSL4 and AR in breast cancer patients. Analyses rendered an inverse expression pattern between these molecules (data not shown). AR and ACSL4 protein expressions were also assessed in breast cancer cell lines. Western blot analyses thus revealed a negative relationship between ACSL4 and AR regardless of the cell model (Fig. 1). It should be noted that AR expression was detected in hormone-positive MCF-7 cells and in MDA-MB-453 cells but not in MDA-MB-231 or BT-20 cells, while ACSL4 expression was observed in MDA-MB-231 and BT-20 cells only. Altogether, these results show an inverse correlation between ACSL4 and AR expression in breast cancer.

**Figure 1.**
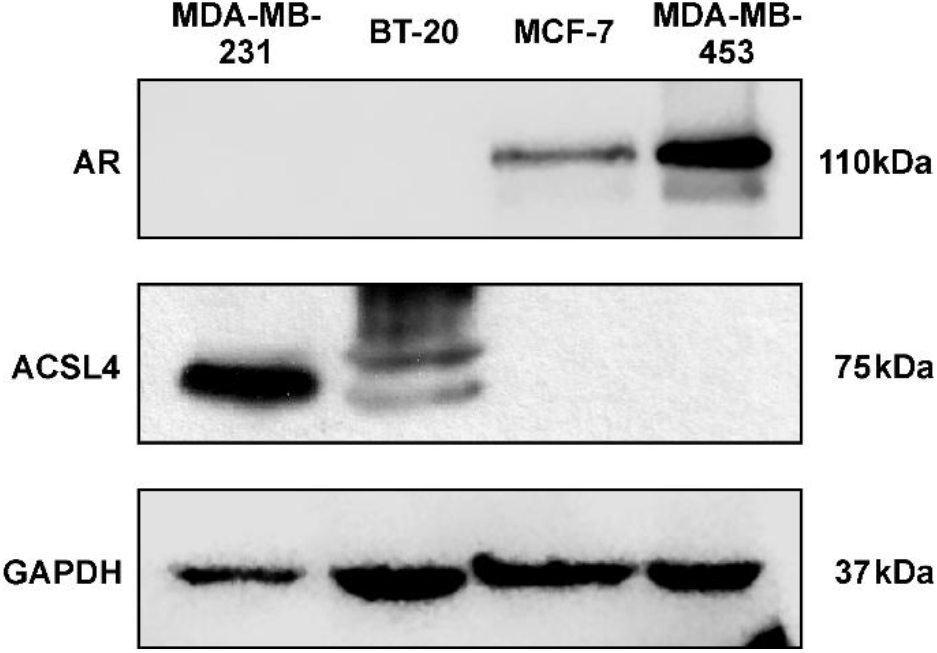
Analysis of ACSL4 and AR expression in breast cancer cells. Protein expression of AR and ACSL4 was analyzed by western blot in total lysates from breast cancer cell lines (MDA-MB-231, BT-20, MCF-7, and MDA-MB-453). GAPDH was used as loading control. A representative blot from at least three independent experiments is shown.

### AR decreases ACSL4 expression in breast cancer cell lines

To determine whether AR regulates ACSL4, we analyzed the effects of ectopic AR expression on ACSL4 levels in breast cancer cells. For this purpose, the TNBC cell lines BT-20 and MDA-MB-231, which express undetectable levels of AR and high levels of ACSL4, were used (Fig. 1). Transient overexpression of AR was induced using a pSG5-ARh plasmid (31), with the empty pSG5 vector used as control. Western blot (Fig. 2 A and B) and qPCR assays (Fig. 2 C and D) showed that ectopic AR in MDA-MB-231 and BT-20 cells led to a decrease in ACSL4 protein and mRNA levels. We next assessed the effect of an antiandrogen on AR-positive, ACSL4-negative MCF-7 cells (Fig. 1) using 48 h incubation with 10 µm BICA. Results showed higher ACSL4 levels in BICA-incubated cells than in those incubated with vehicle alone, as corroborated through immunocytochemistry (Fig. 2 E).

**Fig 2.**
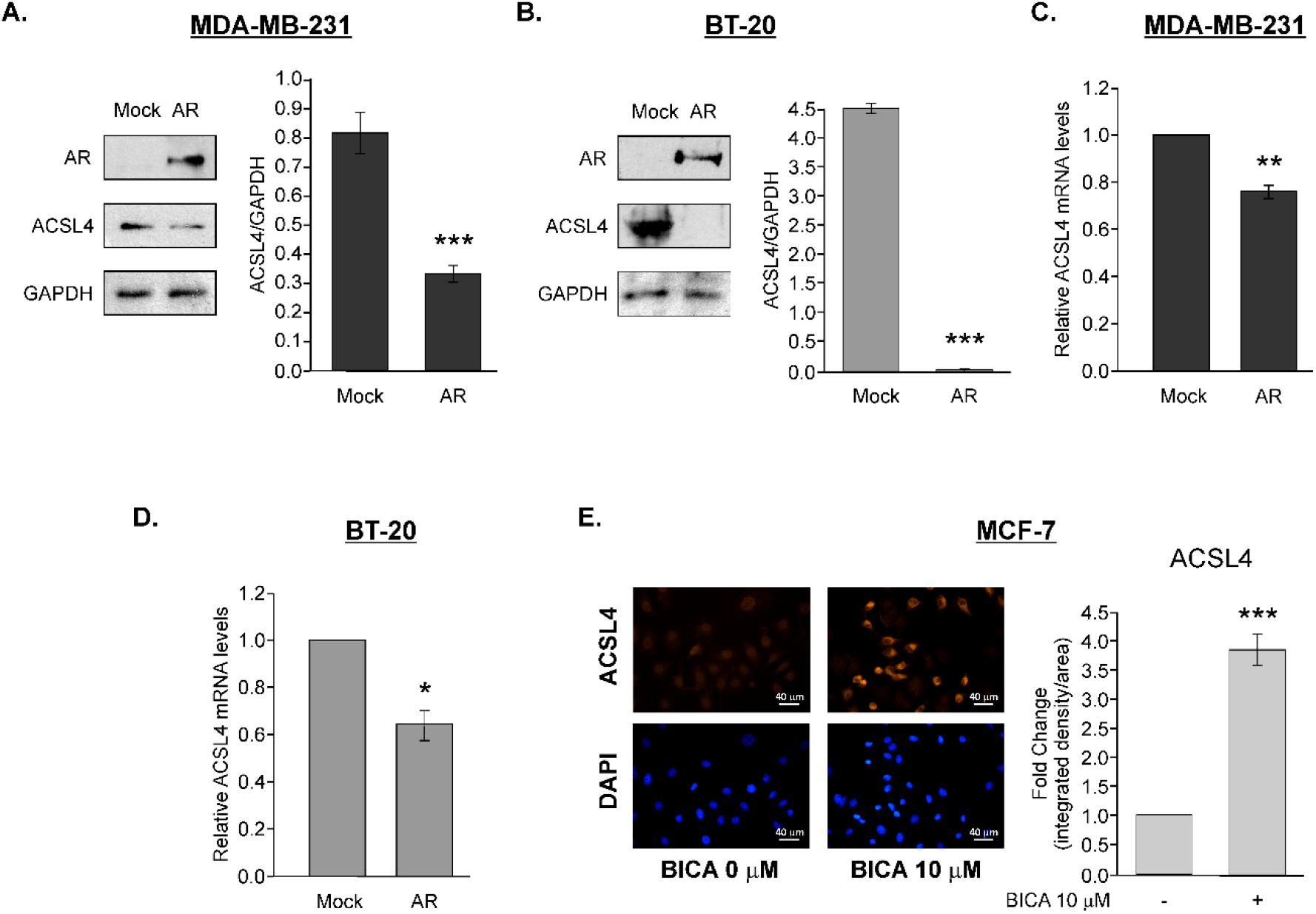
Androgenic regulation of ACSL4 expression in breast cancer cell lines. Effect of ectopic AR expression on ACSL4 protein and mRNA levels. MDA-MB-231 and BT-20 cells were transiently transfected with pSG5-ARh (AR) or empty vector (Mock) and incubated in complete medium for 72 h. **(A–B)** AR and ACSL4 protein expression was analyzed by western blot in total cell lysates. Integrated optical density of ACSL4 was quantified and normalized against GAPDH expression. Representative blots and quantification are shown. Results are expressed as the mean +/- SD in arbitrary units of at least three independent experiments. ***P<0.001 vs. Mock. **(C–D)** ACSL4 mRNA levels were determined by RT-qPCR and normalized against the corresponding from cyclophilin. Results are expressed as the mean +/- SD in arbitrary units of at least three independent experiments. **P<0.01, *P<0.05 vs. Mock. **(E)** Effect of BICA on ACSL4 protein expression in MCF-7 cells analyzed by immunocytochemistry. MCF-7 cells were cultured in complete medium and treated with 10 µM BICA or DMSO as vehicle for 48 h. Representative fluorescence microscopy images at 40X of cells stained with anti-ACSL4 1:100 (red), and nuclei counterstained with DAPI (blue) are shown. Images were obtained at equal exposure time and intensity. Quantification of ACSL4 fluorescence intensity is shown. Control conditions were arbitrarily set to 1. Results are expressed as the mean +/- SD in arbitrary units of at least three independent experiments. ***P<0.001 vs. vehicle.

ACSL4 promoter activity was assessed using the ACSL4 Promoter reporter plasmid which contains a 1.8 kb fragment of the human ACSL4 promoter upstream of the NanoLuc luciferase gene (27). In order to analyze the effect of AR on ACSL4 promoter activity, pSG5-ARh (31) or pSG5 were cotransfected with the ACSL4 promoter in MDA-MB-231 cells. Results showed that AR expression led to a reduction in ACSL4 promoter activity (Fig. 3 A), which is consistent with mRNA and protein analyses. We then studied the effect of AR activation or inhibition on ACSL4 transcription in breast cancer cells. MCF-7 cells were transfected with the ACSL4 Promoter and then treated with AR agonist DHT at a concentration of 10 nM or AR antagonist BICA at a concentration of 10 µM. NanoLuc luciferase reporter assays showed that DHT reduced, whereas BICA increased, ACSL4 transcriptional activity (Fig. 3 B and C). We next aimed to assess whether these effects were specifically due to AR action on the ACSL4 promoter. Therefore, we first performed *in silico* analysis of the human ACSL4 promoter sequence with the ALGGEN PROMO tool. The analysis revealed an ARE present in the 1.8 kb fragment between positions -917 and -899 with a core similarity of 1 and a matrix similarity of 0,925. To further investigate these findings, we performed site-directed mutagenesis of the ARE within the ACSL4 Promoter (Mut ARE) and compared its transcriptional activity to that of the wild-type promoter in MCF-7 cells. The mutation significantly increased ACSL4 promoter activity, which indicates that this ARE functions as a transcriptional repressor (data not shown).

**Fig 3.**
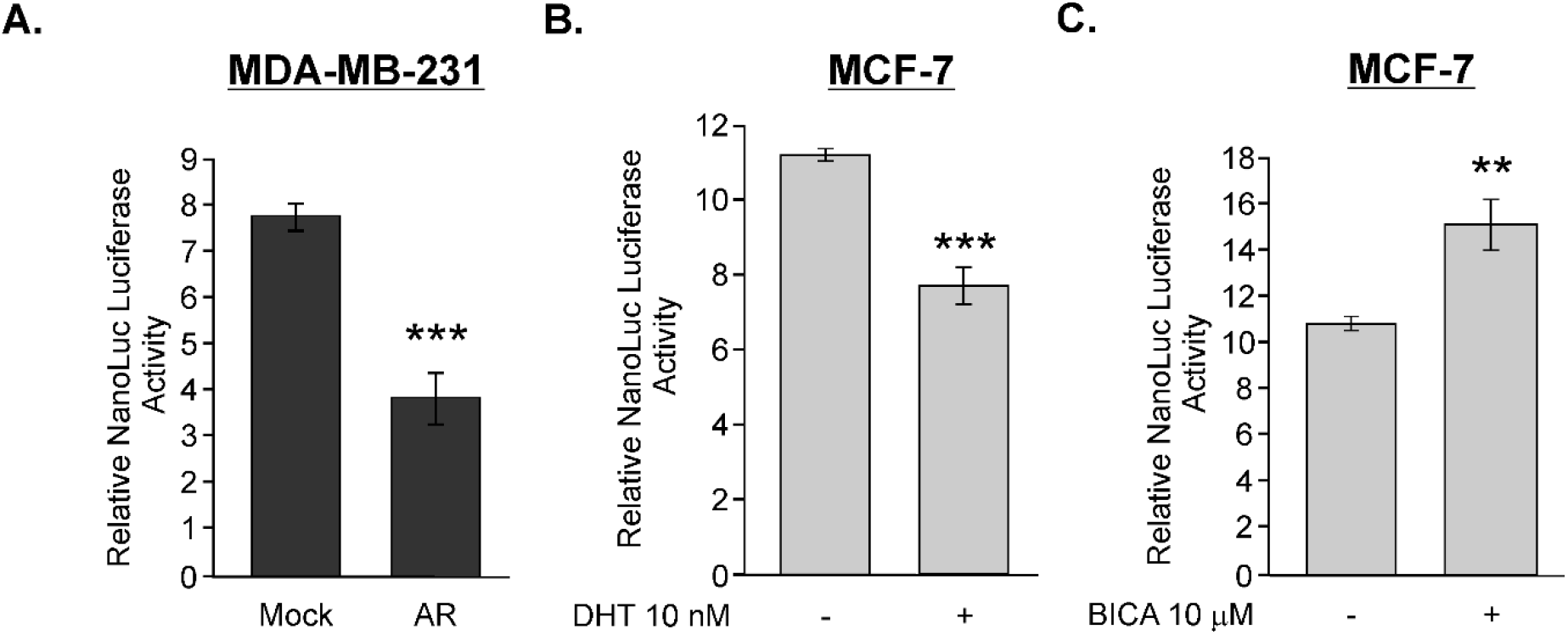
Androgenic regulation of ACSL4 promoter activity in breast cancer cell lines. The construct referred to as “ACSL4 Promoter” consists of a 1.8 kb fragment of the human ACSL4 promoter cloned into the pNL1.1 vector and was used for transfection into breast cancer cells. **(A)** ACSL4 promoter was co-transfected with pSG5-ARh (AR) or empty vector (Mock) as control in MDA-MB-231 cells. Cells were allowed to recover for 72 h and NanoLuc luciferase activity was then determined using the Nano-Glo® Luciferase Assay System (Promega) to measure luminescence. **(B)** Transcriptional activity was measured as previously described in MCF-7 cells transfected with ACSL4 promoter and treated with DHT (10 nM) or vehicle as control for 24 h in steroid-free medium as is described in material and methods. **(C)** Promoter activity was measured as previously described in MCF-7 cells transfected with ACSL4 Promoter and treated with AR antagonist BICA (10 μM) or vehicle as control for 48 h in complete medium. For all experiments, the relative NanoLuc luciferase activity was determined by co-transfection of pRc/CMVi plasmid (Life Technologies) expressing the EGFP gene. Graphics show quantification of arbitrary units of transcriptional activity calculated as luminescence counts related to EGFP fluorescence counts. Results are expressed as the mean +/- SD in arbitrary units of at least three independent experiments. ***P<0.001, **P<0.01 vs. Control.

### ACSL4 negatively modulates AR expression in breast cancer cell lines

Next, we analyzed the effect of ACSL4 overexpression on AR levels. Western blot studies in ACSL4-overexpressing MCF-7 cells revealed a reduction in AR expression upon ACSL4 overexpression (Fig. 4 A). Furthermore, the stable silencing of ACSL4 in MDA-MB-231 cells induced an increase in AR expression (Fig. 4 B). Comparable findings were obtained through ACSL4 inhibition with PRGL493, an ACSL4 inhibitor (24). Treatment with different doses of PRGL493 for 72 h increased AR mRNA levels in MDA-MB-231 cells starting at a PRGL493 concentration of 2.5 µM (Fig. 4 C) and increased AR protein expression in both MDA-MB-231 and BT-20 cells as from a PRGL493 concentration of 10 µM (Fig. 4 D and E). Overall, these results indicate that inhibition of ACSL4 induces AR expression in AR-negative MDA-MB-231 and BT-20 breast cancer cells.

**Figure 4.**
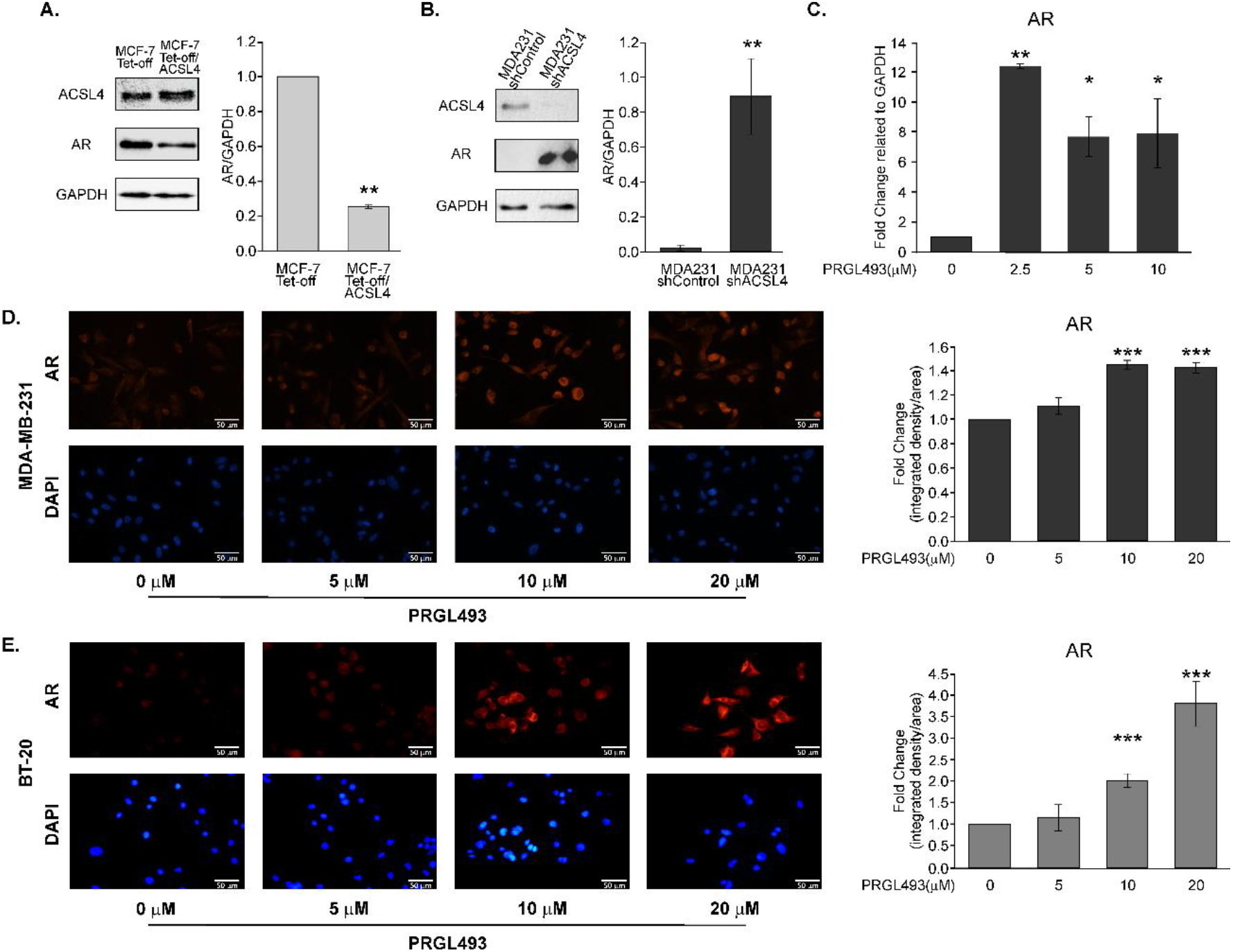
ACSL4 modulation regulates AR expression in breast cancer cells. Western blot analysis of AR and ACSL4 protein expression in **(A)** stable MCF-7 Tet-Off/ACSL4 or MCF-7 Tet-Off cells as control or in **(B)** stable MDA-MB-231 shACSL4 and MDA-MB-231 shControl cells as control. AR levels were quantified by densitometry and normalized to GAPDH. Representative blots and densitometric quantification are shown. **(C)** AR mRNA expression was determined by RT-qPCR in cells treated with increasing concentrations of PRGL493 (0–10 µM) for 72 h in complete medium. mRNA levels were normalized to GAPDH and expressed as fold change relative to control (cells incubated with vehicle). Effect of PRGL493 treatment (0–20 µM, 72 h) on AR protein expression in **(D)** MDA-MB-231 and **(E)** BT-20 cells analyzed by immunocytochemistry. Representative fluorescence microscopy images at 40X of cells were stained with anti-AR 1:100 (red), and nuclei were counterstained with DAPI (blue) are shown. Cells incubated with vehicle were used as control. Images were obtained at equal exposure time and intensity. Quantification of AR immunofluorescence is shown on the right. Results are expressed as the mean ± SD of at least three independent experiments vs. corresponding control. ***P<0.001, **P<0.01, *P<0.05 vs. Control.

### The combination of PRGL493 and BICA reduces cell aggressiveness in breast cancer

We have previously shown that ACSL4 inhibition with PGRL493 sensitizes MDA-MB 231 cells to chemotherapy or hormonal therapy, promoting the expression of ER (24). Similar results have been reported in studies on MDA-MB-231 and prostate cancer cells PC-3, in which PRGL493 was combined with chemotherapeutic agents (24). Furthermore, understanding the molecular mechanisms underlying ACSL4 regulation and its contribution to tumor development and progression may provide valuable insights for the development of effective therapeutic strategies, including those aimed at modulating ACSL4 expression (13,15,24). As we observed that ACSL4 inhibition with PRGL493 increased AR levels in cells lacking this receptor, we next investigated whether ACSL4 inhibition may sensitize breast cancer cells to antiandrogens using combined pharmacological approaches involving submaximal doses of PRGL493 and BICA. A dose-response cell proliferation curve was initially generated for both inhibitors. Results showed PRGL493 inhibitory effects on cell proliferation starting at a concentration of 4 µM but no inhibitory effects for BICA up to a concentration of 20 µM (Fig. 5 A). For combination assays, MDA-MB-231 cells were pre-incubated with PRGL493 for 48 h and then incubated with the combination of BICA and PRGL493 for another 48 h. Results showed synergistic inhibition of MDA-MB-231 proliferation when 3 or 4 µM BICA was combined with 2 or 2.5 µM PRGL493. A strong and consistent synergy was demonstrated in all combined BICA and PRGL493 treatments according to the Bliss independence model (40), with 4 µM BICA plus 2.5 µM PRGL493 standing out as the most effective combination with an EOB of 39.4% (Fig. 5 B). Wound healing assays were then performed to evaluate the effectiveness of combined treatment in curbing cell migration. A heightened inhibitory effect on migration with respect to the inhibitors used alone was observed when MDA-MB-231 cells were exposed to 4 µM BICA and 2.5 µM PRGL493. This combined treatment exhibited synergistic effects, as the difference between the inhibition observed and expected yielded an EOB of 14% in Bliss analysis. Worth pointing out, the migration in cells submitted to single treatment did not differ significantly from that of cells incubated with vehicle alone (Fig. 5 C). The 3D spheroid invasion assay into surrounding Matrigel® was then performed to assess the capacity of tumor cells to invade a surrounding matrix. Initially, a dose-response curve for PRGL493 was determined to ascertain inhibitor effects on the invasion of MDA-MB-231 cells from spheroids, with results showing a significant decrease in cell invasion as from a PRGL493 dose of 10 µM (Fig. 5 D). Subsequent drug combination analyses rendered results consistent with those of the previous proliferation and migration assays. Invasion was similar to that of control cells when 4 µM BICA or 2.5 µM PRGL493 were used alone but suppressed upon combined treatment. Once again, we found a strong synergy in the combination of sub-effective doses of the drugs, as verified by Bliss analysis (44%) (Fig. 5 E).

**Fig 5.**
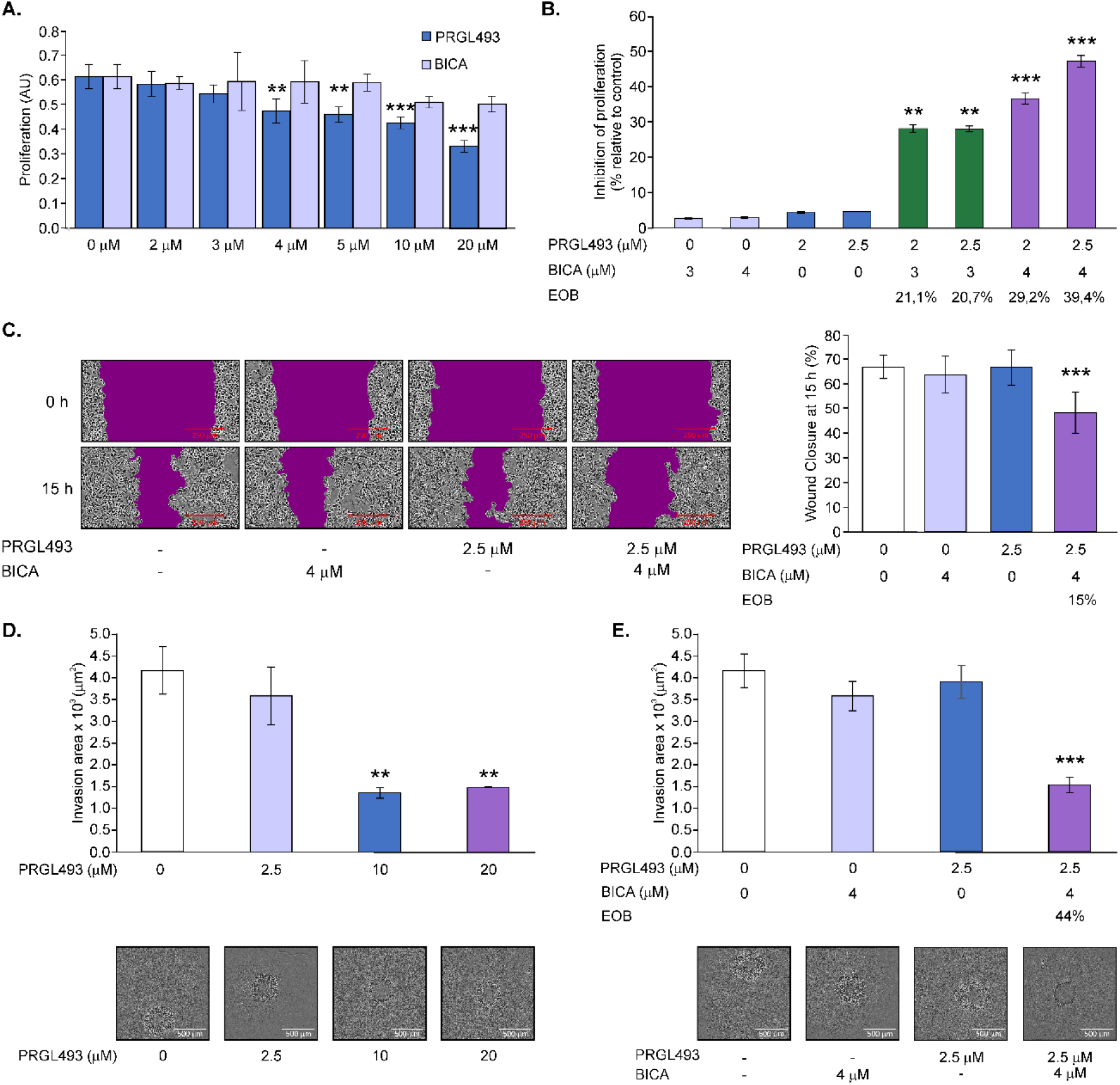
ACSL4 inhibitor PRGL493 sensitizes MDA-MB-231 cells to AR antagonist BICA. **(A)** Dose–response effect of the ACSL4 inhibitor PRGL493 and AR antagonist BICA on the proliferation of MDA-MB-231 cells. Cell proliferation was determined by MTT assay after treatment with increasing concentrations of each compound (0–20 µM) for 96 h. Cells incubated with vehicle alone were used as control. (**p < 0.01; ***p < 0.001 vs. control). **(B)** Effect of combined treatment with PRGL493 and BICA on cell proliferation. Except for cells treated with BICA alone at 3 µM or 4 µM, cells were pre-incubated with PRGL493 2 µM or 2.5 µM for 48 h followed by 48 h incubation with PRGL493 at 2 µM or 2.5 µM and/or BICA at 3 µM or 4 µM. Proliferation inhibition is shown as percentage relative to control cells, which were incubated with vehicle for 96 h. Excess over Bliss (EOB) values are indicated for the combinations (**p < 0.01; ***p < 0.001 vs. corresponding single treatments). **(C)** Effect of combined treatment with PRGL493 and BICA on cell migration assessed through the wound-healing assay in MDA-MB-231 cells. Wounds were made and cells were incubated with PRGL493 (2.5 µM) and/or BICA (4 µM) for 15 h in complete medium. Representative images at 0 h and 15 h acquired using the IncuCyte Live-Cell Analysis System at 10X magnification are shown. The graph displays the percentage of wound closure for each treatment condition at 15 h with respect to 0 h and the corresponding EOB value for the combination. (***p < 0.001 vs. corresponding single treatments) **(D)** Dose–response effect of the ACSL4 inhibitor PRGL493 on the invasion of MDA-MB-231 cells. Cell invasion was determined by 3D Spheroid Invasion Assay after treatment with increasing concentrations of PRGL493 (0–20 µM) in complete medium and 25% of matrigel for 5 days. Cells incubated with vehicle alone were used as control. Representative images on day 5 acquired using the IncuCyte Live-Cell Analysis System at 4X magnification are shown. The upper graph shows the cell invasion area with respect to the spheroid core for each treatment condition on day 5. (**p < 0.01 vs. control). **(E)** Effect of ACSL4 inhibitor PRGL493 and/or AR antagonist BICA on the invasion of MDA-MB-231 cells. Cell invasion was assessed by 3D Spheroid Invasion Assay after treatment withPRGL493 (2.5 μM) and/or BICA (4 μM) in complete medium and 25% of matrigel for 5 days. Cells incubated with vehicle alone were used as control. Representative images on day 5 acquired using the IncuCyte Live-Cell Analysis System at 4X magnification are shown. The upper graph shows the cell invasion area with respect the spheroid core for each treatment condition on day 5 and the corresponding EOB value for the combination (***p < 0.001 vs. corresponding single treatments). Results are expressed as the mean ± SD of at least three independent experiments.

### The combination of PRGL493 and BICA suppresses extravasation in breast cancer cells

Finally, we tested the effect of PRGL493 and BICA on MDA-MB-231 cell extravasation, a key step in metastasis, using the zebrafish model. Prior to xenotransplantation, cells were incubated with different doses of PRGL493. Live imaging on a confocal microscope was then used to track cell migration through the vascular system and extravasation after 24 h. Results showed the inhibition of cancer cell extravasation in zebrafish at a PRGL493 concentration of at least 2.5 µM (Fig. 6 A), which provides evidence of the biological effects of pharmacological ACSL4 inhibition in this crucial metastasis step. Subsequently, we tested the effect of drug combinations on cancer cell extravasation. As cells incubated with 1.5 µM PRGL493 showed widespread extravasation in the dose-response curve (Fig. 6 A), we chose this concentration for combined treatment. Before intracardiac microinjection, except for the group injected with 4 µM BICA alone and the control group, cells were pre-incubated with 1.5 µM PRGL493 for 24 h and then incubated with 4 µM BICA and/or 1.5 µM PRGL493 for another 48 h. MDA-MB-231 cell incubation with the combination of drugs suppressed extravasation, as different from single treatment or vehicle alone (Fig. 6 B). These results demonstrate the efficacy of combined submaximal doses of PRGL493 and BICA in suppressing the behavior of cancer cell in extravasation *in vivo*.

**Fig 6.**
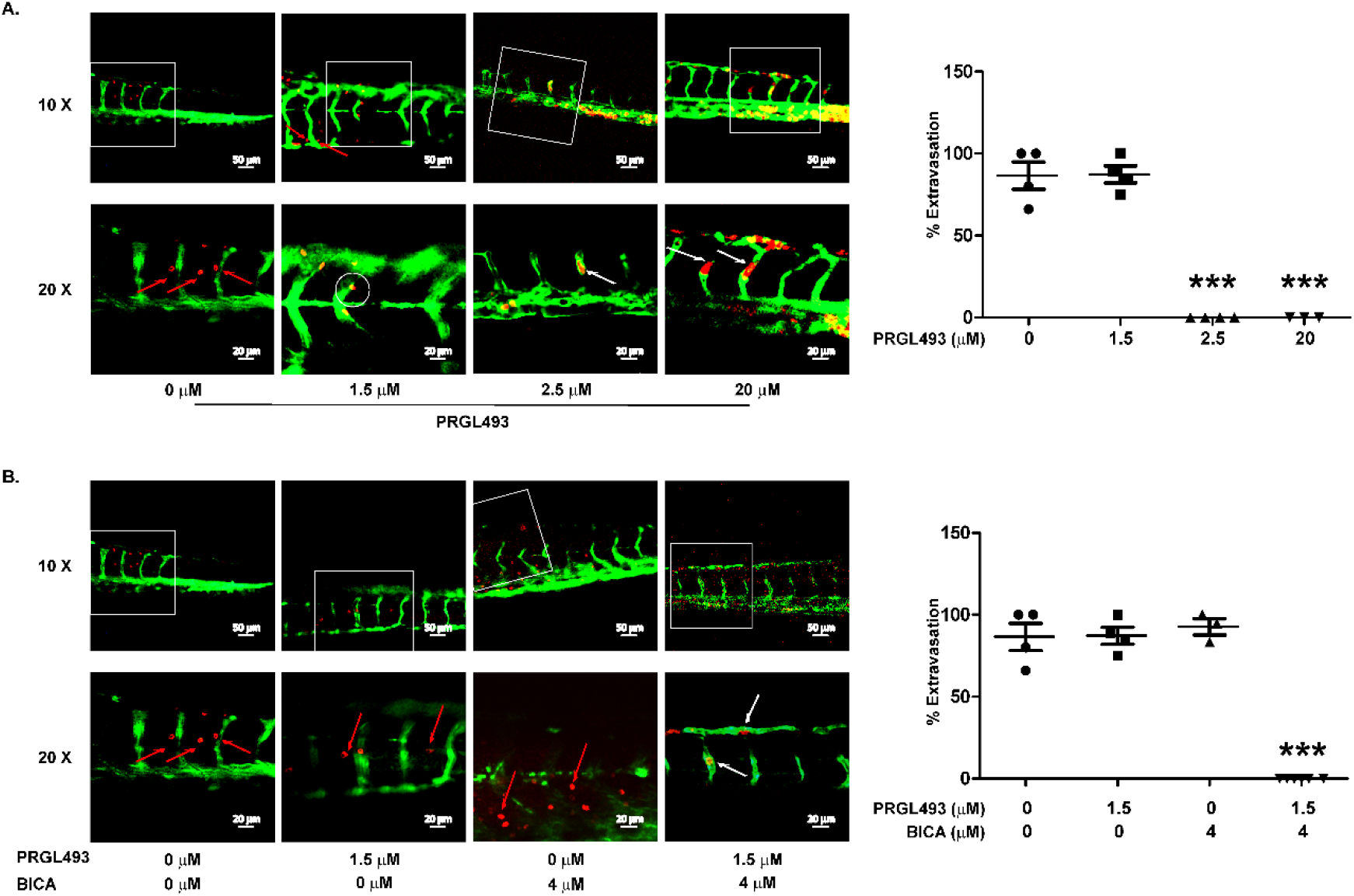
The ACSL4 inhibitor PRGL493 suppresses MDA-MB-231 extravasation in a zebrafish xenograft model both alone and in combination with BICA. **(A)** Effect of increasing concentrations of PRGL493 (0–20 µM) on MDA-MB-231 breast cancer cell extravasation in a zebrafish model. Cells previously incubated with the indicated dose of PRGL493 for 72 h were labeled with a red fluorescent tracker dye and then microinjected intracardiacally in 3-dpf *Tg(fli-GFP)* zebrafish embryos with the endothelium labeled with the green EGFP protein reporter. **(B)** Effect of combined treatment with PRGL493 (1.5 µM) and BICA (4 µM) on MDA-MB-231 cell extravasation in a zebrafish model. Except for cells treated with BICA alone (4 µM), cells were pre-incubated with PRGL493 (1.5 µM) for 24 h followed by 48 h incubation with PRGL493 1.5 µM and/or BICA 4 µM. Cells without treatment were incubated with vehicle for 72 h. Once the cell incubation period was over, cells were labeled with a fluorescent tracker dye (red) and microinjected intracardiacally in 3-dpf *Tg(fli-GFP)* zebrafish embryos with endothelium labeled with the green EGFP protein reporter. All images were acquired 24 h after xenotransplantation using the Zeiss LSM 900 confocal microscope and the photographs were acquired using the ZEISS ZEN 3.11 software. Representative images at 10X and 20X magnification are shown. The 20X images are a zoom of the area framed in the corresponding 10X image. White arrows point out the cells remaining in circulation within the vessels, without extravasation. The circle shows a cell actively extravasating. Red arrows show examples of cells that have already extravasated. The graphs on the right show the percentage of extravasation in zebrafish embryos. Results are expressed as the mean ± SD of at least three independent embryos vs. corresponding control. ***P<0.001 vs. cell without treatment.

## Discussion

In this study, we demonstrate a novel therapeutic strategy for aggressive breast cancer by establishing and exploiting a reciprocal negative regulation between AR and ACSL4. Our findings reveal that the pharmacological inhibition of ACSL4 not only upregulates AR expression but also critically sensitizes classically AR-negative breast cancer cells to antiandrogen therapy. Furthermore, we establish that a dual metabolic and endocrine blockage using submaximal doses of ACSL4 and AR inhibitors synergistically suppresses tumor cell extravasation. Finally, this work provides a strong mechanistic rationale for expanding AR-targeted therapies to QNBC patients, addressing a critical unmet clinical need.

Mechanistically, we first established that AR diminishes ACSL4 levels in breast cancer. ACSL4 expression or promoter activity were reduced by AR overexpression or activation via agonist DHT but increased by AR antagonist BICA. Moreover, site-directed mutagenesis of the ARE in the ACSL4 promoter confirmed AR’s role as ACSL4 repressor. Notably, the regulatory dynamics observed in our breast cancer models are consistent with mechanisms previously documented in prostate cancer (22,28,41). In addition, we demonstrate that ACSL4 negatively regulates AR expression in breast cancer cells. While previous reports showed that ectopic ACSL4 decreases AR mRNA in those models (25), we determined that it also reduces AR protein levels. Most importantly, the silencing or pharmacological inhibition of ACSL4 with PRGL493 significantly increased AR expression at both the protein and mRNA levels. This reciprocal feedback loop highlights a metabolic-endocrine crosstalk that can be therapeutically leveraged.

Building on this mechanistic link, we evaluated the therapeutic potential of upregulating AR via ACSL4 inhibition in TNBC. While AR is a potential therapeutic target in AR-positive TNBC subsets (4,10), benefits remain restricted to tumors with detectable AR expression. We thus hypothesized that ACSL4 inhibition could sensitize AR-low or AR-negative cells to antiandrogens, in line with approaches reversing tamoxifen resistance in ER-positive models (23,24). Our results confirmed that combined treatment using PRGL493 and BICA significantly reduced proliferation, migration, and invasion in MDA-MB-231 cells. Even though the AR status of MDA-MB-231 cells is debated —often considered a QNBC phenotype due to the lack of baseline AR expression (25,42–44)—, the fact that PRGL493 successfully sensitized these aggressive cells to BICA demonstrates that metabolic modulation can overcome intrinsic endocrine resistance.

Metastasis is a multistep process involving local invasion, intravasation into blood or lymphatic vessels, survival in the circulation, extravasation from blood vessels into distant tissues, and colonization, which ultimately leads to the formation of secondary tumors (45). Given that metastasis is the primary cause of breast cancer mortality, we also assessed cancer cell extravasation, a critical step in the metastatic process (46). ACSL4 has been documented as a metastasis-inducing factor (47), previously linked to cancer cell extravasation in ovarian and liver cancers (48). We used the transgenic zebrafish strain *Tg(fli1:EGFP)* to evaluate its role in breast cancer cell extravasation. We demonstrate that PRGL493 alone profoundly suppressed MDA-MB-231 cell extravasation at concentrations of at least 2.5 µM. The dual pre-treatment with PRGL493 and BICA have changed the subsequent behaviour of the cells after injection, suppressing cancer cell extravasation at submaximal concentrations that showed no efficacy in single-agent treatments.

The robust effects observed in our models underscore the potential of PRGL493 as a novel, clinically viable therapeutic agent. The pharmacological targeting of ACSL4 using PRGL493 has demonstrated efficacy in prostate and breast cancer (24,49). Furthermore, the therapeutic scope of ACSL4 inhibition by PRGL493 is expanding into other pathological contexts characterized by lipid metabolism dysregulation (50–52). By establishing that PRGL493 can be effectively combined with endocrine therapies at submaximal doses, our study not only reinforces the potential of PRGL493 as possible therapeutic agent but also supports that combined ACSL4/AR inhibition has potential to represent a promising therapeutic strategy to impair cancer cell extravasation, a key step of metastatic dissemination. These findings provide a preclinical framework for its future translation into clinical trials since further studies are needed to determine whether this strategy also affects other steps of the metastatic cascade and study whether the strategy can effectively limit metastatic progression and to support its potential clinical translation.

QNBC presents an urgent clinical challenge characterized by early metastasis, limited treatment options, and poor prognosis. Here, by demonstrating that ACSL4 suppresses AR expression, we have identified a metabolic switch which can be therapeutically targeted to functionally reprogram AR-negative tumors to respond to antiandrogens. The synergistic efficacy of combined ACSL4 and AR inhibition in halting cancer aggressiveness highlights a promising translational approach. Combination strategies targeting metabolic vulnerabilities are increasingly recognized for overcoming resistance and improving tolerability (53,54). In this context, our proposed dual intervention does not only maximize antitumor efficacy but also minimizes potential systemic toxicity. These results thus pave the way for innovative clinical trials that combine metabolic modulators and endocrine therapies to broaden the clinical applicability of AR-directed treatments in aggressive breast cancers.

